# General encoding of canonical *k*-mers

**DOI:** 10.1101/2023.03.09.531845

**Authors:** Roland Wittler

## Abstract

To index or compare sequences efficiently, often *k*-mers, i.e., substrings of fixed length *k*, are used. For efficient indexing or storage, *k*-mers are often encoded as integers, e.g., applying some bijective mapping between all possible *σ*^*k*^ *k*-mers and the interval [0, *σ*^*k*^ −1], where *σ* is the alphabet size.

In many applications, e.g., when the reading direction of a DNA-sequence is ambiguous, *canonical k*-mers are considered, i.e., the lexicographically smaller of a given *k*-mer and its reverse (or reverse complement) is chosen as a representative. In naive encodings, canonical *k*-mers are not evenly distributed within the interval [0, *σ*^*k*^ −1].

We present a minimal encoding of canonical *k*-mers on alphabets of arbitrary size, i.e., a mapping to the interval [0, *σ*^*k*^*/*2−1]. The approach is introduced for canonicalization under reversal and extended to canonicalization under reverse complementation. We further present a space and time efficient bit-based implementation for the DNA alphabet.

## 1 Introduction

The increasing amount of available genome sequence data enables large-scale analyses. A common way to handle long and many genomes is using *k*-mers, i.e., substrings of fixed length *k*, to efficiently index or compare sequences.

Many *k*-mer-based methods hash *k*-mers to integer values in order to store them in a table, or to assign them to different tables or threads.

For a specific subset *S* of observed *k*-mers among all possible *k*-mers, *minimal perfect hash functions* (MPHFs) can be used. Here, a data structure comprising the given set *S* is used to bijectively map an individual *k*-mer to an integer (hash value) in the interval [0, |*S*|−1], see e.g., [2, 3, 5, 6, 7, 8].

In contrast, here, we consider a general encoding of any *k*-mer from the universal set of *k*-mers that is independent of any data set but may be restricted by certain properties as described below.

For the four letter DNA alphabet, each character can be encoded using two bits. Reading the resulting bit sequences as integer values corresponds to an encoding for the set of all 4^*k*^ *k*-mers which are mapped to an interval of size 4^*k*^.

In many situations, e.g., when genomes are not given as complete sequences but rather in form of contigs or reads, the reading direction of a given DNA sequence is unknown. To make a method independent from whether a sequence itself of its reverse (Watson Crick) complement is observed, both a *k*-mer and its reverse complement are assumed equivalent and one of them is chosen as a *canonical* representative of both, e.g., the lexicographically smaller one. For odd length *k*-mers, two *k*-mers pair to one canonical *k*-mer each. Thus, there are half as many canonical *k*-mers as there are *k*-mers. For even length *k*-mers, (Watson Crick) palindromes have to be considered, which are canonical by definition and do not pair, thus increasing the number of canonical *k*-mers. In any case, the (lexicographically) canonical *k*-mers are not evenly distributed – neither within the lexicographic ordering of *k*-mers nor in the interval of *k*-mer hash values spanned by a simple 2-bit encoding. See Figure 1 for an illustration.

**Figure 1:**
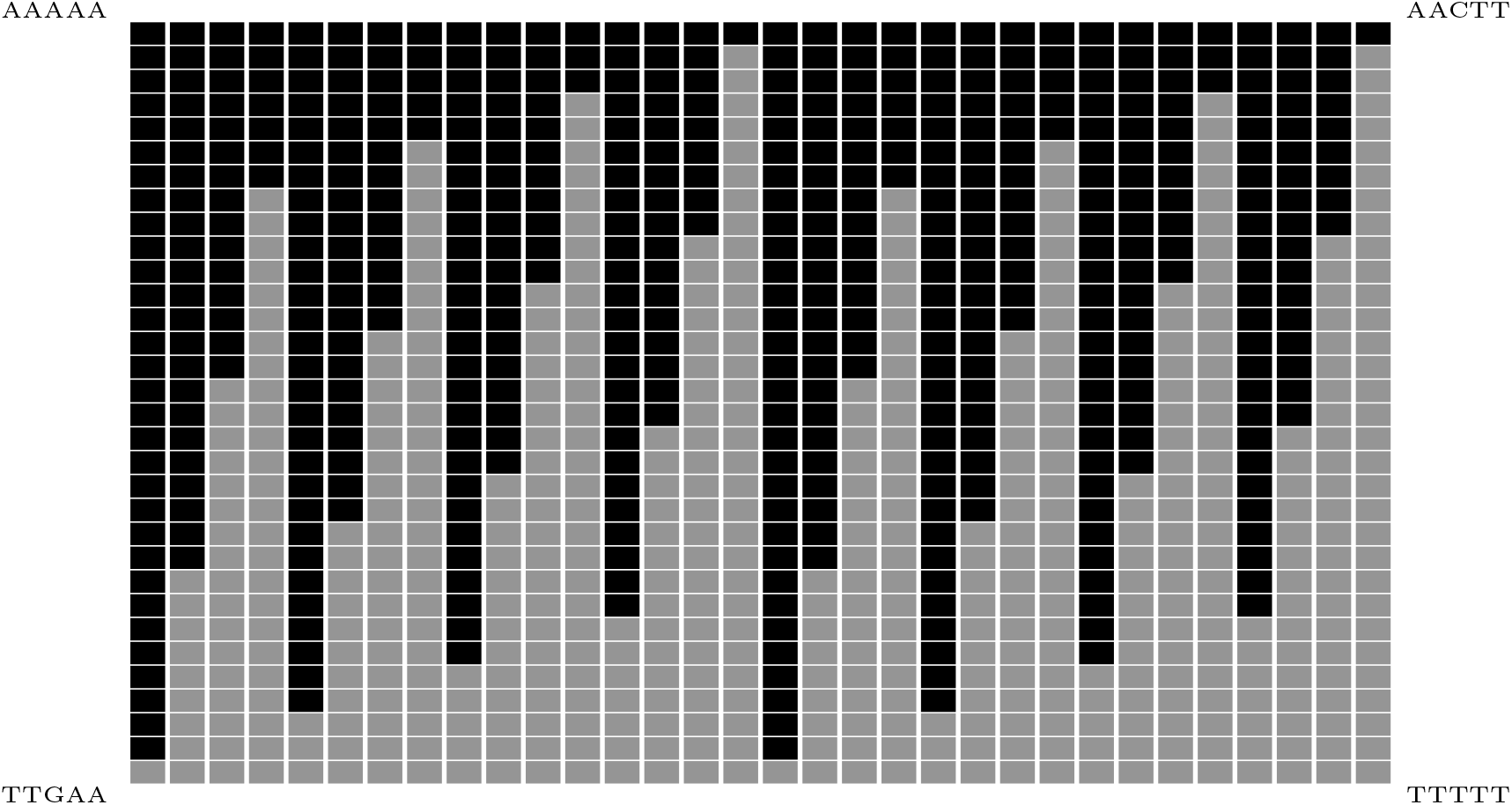
Distribution of (lexicographically) canonical *k*-mers among all *k*-mers. Each *k*-mer for *k* = 5 is represented as a block, ordered lexicographically from left to right, row by row. The four blocks at the corners are exemplarily labeled. A block is drawn black if the *k*-mer is canonical, and gray otherwise.

The canonical *k*-mers correspond to only a subset of all hash values. In view of the fact that (for odd *k*) exactly half of all *k*-mers are canonical, only 2*k* − 1 bits instead of 2*k* bits are necessary and sufficient to span the integer values from 0 to *n* −1 where *n* is the number of canonical *k*-mers. One approach to obtain a 2*k* −1-bit encoding is to define a *k*-mer being canonical not with respect to their lexicographic relation but rather based on their 2-bit encoding. To the best of our knowledge, such an approach has not been published yet, but has been proposed by the community^1^: Encode characters by two bits in such a way that a character and its complement have different parity each (even or odd number of ones), e.g., *A* ↦ 00, *C* ↦ 01, *G* ↦11, *T* ↦ 10. Then, the encoding of an odd length *k*-mer always has a different parity as its reverse complement. Hence, one parity can be chosen as canonical, and one bit, e.g. the first bit, can be omitted as it could be deduced by counting the ones in the shortened bit sequence. This approach, however, is only applicable for odd values of *k*, and there is no evident extension to even length *k*-mers.

While all above considerations are based on the DNA alphabet of size four, also alphabets of other sizes are worth to study. An alphabet size *σ* that is not a power of two does not allow for simple bit-based encoding. A generalization to a ranking-based approach offers an elegant integer encoding: Interpret a character as a digit between 0 and *σ* − 1, and a *k*-mer as a number in a base-*σ* numeral system. Then, a standard mapping to decimal values corresponds to an encoding to [0, *σ*^*k*^ −1].

We stress that, on the one hand, we are not aware of any case where the issues outlined above lead to serious limitations in practice. Exact encodings are usually not necessary in practice, randomized hashing is applied [1, 4, 9] and even-length *k*-mers are often omitted anyway to avoid palindromes. On the other hand, we observe an interesting theoretical phenomenon: while encoding, enumerating etc. *k*-mers is straight-forward and also the definition of being *canonical* is simple, in contrast, encoding canonical *k*-mers appears to be complex. We want to contribute to the basic research in this field by filling this gap in the theory of *k*-mers. We propose a general, minimal encoding for canonical *k*-mers – *general* in the sense that it applies for arbitrary *k* (even or odd) and on alphabets of arbitrary size *σ*, and *minimal* in the sense that it bijectively maps all *n k*-mers to the interval [0, *n* −1]. The approach is introduced for canonicalization under reversal and extended to canonicalization under reverse complementation. For alphabet size four, we show the relation to bit-space and provide an algorithm to encode all *k*-mers of a given sequence *s* in time *O*(*k* + |*s*|). A simple implementation is available on GitLab: https://gitlab.ub.uni-bielefeld.de/gi/MinEncCanKmer

## 2 Preliminaries

We consider as *alphabet* a finite, non-empty, ordered set {*a*_0_, … , *a*_*σ*−1_} with *a*_0_ *<* … *< a*_*σ*−1_ and define the *rank* of character *a*_*i*_ ∈ 𝒜 as *rank*(*a*_*i*_) = *i*. A *string s* is a sequence of characters. Its *length* is denoted by |*s*|, the character at position *i* by *s*[*i*], and the substring from position *i* through *j* by *s*[*i*..*j*], where 1 ≤*I* ≤ *j*≤ |*s*|. Let *ϵ* denote the empty string. The concatenation of a string or character *s* and another string or character *t* is denoted by *s* ·*t*. The *reverse* of string *s* is *s*^−1^ = *s*[|*s*|] … *s*[1].

Strings of fixed length *k* are called *k-mer*. We define a *k*-mer *x* being *canonical* if it is lexicographically smaller than or equal to *x*^−1^.

Let 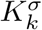 be the set of all *k*-mers over an alphabet of size *σ*. The number of *k*-mers is 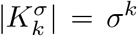. For the number of canonical *k*-mers, first note that a *palindrome*, i.e., a sequence *x* = *x*^−1^, is canonical by definition. Each non-palindromic *k*-mer pairs with its reverse such that the number of canonical non-palindromic *k*-mers is half of the number of non-palindromic *k*-mers. The number of palindromes is 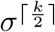, so the number of non-palindromic *k*-mers is 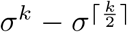, half of which are canonical. In summary, the size of the set 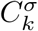 of canonical *k*-mers is:

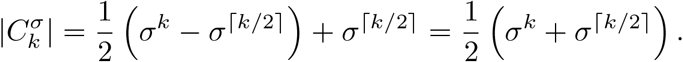

In order to efficiently handle *k*-mers, they can be mapped to integers.

### Definition 1.

*An* encoding *on U is an injective total function that maps elements from U to integer values. It is* minimal *if it bijectively maps to the range* [0, |*U* | − 1].

Please note the following:

- In other contexts, encodings can be defined to map on words over specific alphabets. Here we assume mappings to single integers.
- We use the term *encoding* for both, a numeral representing a *k*-mer, also called *code word*, as well as the mapping process itself.
- While formally, the notion of a *minimal encoding* is equivalent to a *minimal perfect hash function (MPHF)*, they are commonly used in different settings: the first on an entire universe of elements, e.g., all possible *k*-mers, and the latter on a given subset of elements, e.g., all *k*-mers observed in a data set. Here, we address minimal encodings of *k*-mers.

A commonly used minimal encoding of *k*-mers is rank based. Each character is encoded by its rank, which is interpreted as a digit. The resulting sequence of digits can then be converted from the base-*σ* numeral system, denoted by (… )_*σ*_, to decimal values.

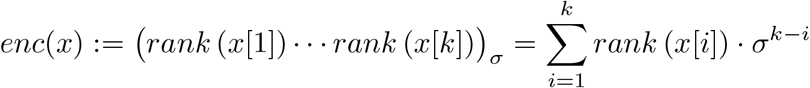

The resulting values range from *enc*(*a*_0_ … *a*_0_) = 0 to 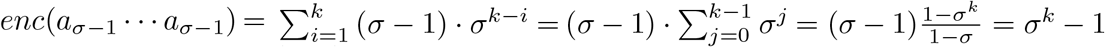, i.e., from 0 to 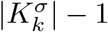.

When instead canonical *k*-mers are encoded using this approach, only part of the co-domain is actually used. In particular, the image does not correspond to an interval of integers. The following section introduces a minimal encoding of canonical *k*-mers, i.e., it has as co-domain the integer interval 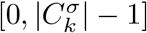.

## 3 Encoding canonical *k*-mers

In the following, we will first introduce a preliminary encoding 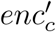 of canonical *k*-mers, which is not minimal. In a second step, gaps in the image of 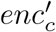 will be removed to derive a minimal encoding *enc*_*c*_. Finally, an expedient characteristic of *enc*_*c*_ with regard to palindromes will be highlighted.

### 3.1 Preliminary encoding

The encoding is based on the observation that there are fewer canonical *k*-mers than there are *k*-mers in general, i.e. 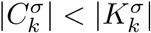. A mapping is only required if *k*-mer *x* is canonical, i.e., *x* is lexicographically smaller than or equal to *x*^−1^. This in turn can be seen when comparing its first to its last character, and if they are equal, its second to its second last etc., i.e., *x*[*i*] to *x*^−1^[*i*] = *x*[*k* − *i* + 1] for *i* = 1, … , *k/*2 until a pair 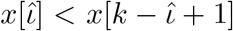 is found, specifying *x* being canonical. If instead equality holds for all pairs, *x* is palindromic and thus canonical by definition.

A given *k*-mer will be encoded from the outside inwards, i.e. for increasing *i*, while different encoding schemes are used. For sake of convenience, the encoding is given as digits in base-*σ* space.

#### Unspecific pairs

If *x*[*i*] = *x*[*k* − *i*+1], one character implies the other such that only one of the two characters needs to be encoded. Instead of actually encoding *x*[*i*], digit zero is used to indicate the pair being unspecific, and *x*[*k* − *i* + 1] is encoded by its rank.

#### Specifying pair

If *x* is not palindromic, let 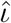 be the smallest value such that 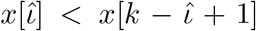. This pair determines *x* being canonical. Only a limited number of combinations of characters needs to be encoded. The left character has to be encoded with a digit greater than zero to tell it apart from the unspecific case. The right character is encoded with a digit from zero to *σ* −1. The combination of digits has to be continuous to avoid gaps in the co-domain. For instance, for an alphabet of size 4, six different combinations of characters are encoded by the lexicographically lowest six pairs of digits from (1, 0) to (2, 1). Function *R* is used to determine the rank of a character pair in the list of all possible combinations, see Table 1 (left) for an example. Values for *R*(*l, r*) can be determined using the *triangular numbers T* = [1, 3, 6, 10, 15, …] (sequence A000217 in the On-Line Encyclopedia of Integer Sequences) as demonstrated in Table 1 (right). The value of *R* is then decomposed into the encoding *R*_*l*_ of the left character and *R*_*r*_ of right character using integer division and modulo, respectively.

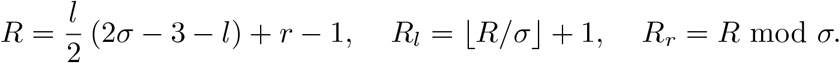

**Table 1:** Example on the encoding of specifying pairs for alphabet size *σ* = 4. The six combinations of characters are listed in lexicographic order and numbered using function *R*. This value is then decomposed to obtain the lexicographically smallest six combinations of digits that do not have a zero in its first position. The tables on the right demonstrate the relation between the *triangular numbers T* = [1, 3, 6, …] (sequence A000217 in the On-Line Encyclopedia of Integer Sequences), highlighted in bold face, and the rank *R*. In the modified matrix *T*_*σ*−1_ − *R*(*l, r*), the values next to the diagonal correspond to the triangular numbers *T*_*σ*−1_, … , *T*_1_. From those, the off-diagonal values can then be easily deduced.

| specifying pair | | rank | encoding | | $T_3 - R$ | 0 | 1 | 2 | 3 |
| --- | --- | --- | --- | --- | --- | --- | --- | --- | --- |
| $l$ | $r$ | $R$ | $R_l$ | $R_r$ | 0 | | **6** | 5 | 4 |
| $a_0$ | $a_1$ | 0 | 1 | 0 | 1 | | | **3** | 2 |
| $a_0$ | $a_2$ | 1 | 1 | 1 | 2 | | | | **1** |
| $a_0$ | $a_3$ | 2 | 1 | 2 | 3 | | | | |
| $a_1$ | $a_2$ | 3 | 1 | 3 | $R$ | 0 | 1 | 2 | 3 |
| $a_1$ | $a_3$ | 4 | 2 | 0 | 0 | | 0 | 1 | 2 |
| $a_2$ | $a_3$ | 5 | 2 | 1 | 1 | | | 3 | 4 |
|  |  |  |  |  | 2 |  |  |  | 5 |
|  |  |  |  |  | 3 |  |  |  |  |

#### Remainder

All remaining characters for 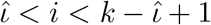 (if any) are simply encoded by their rank.

Our preliminary encoding of canonical *k*-mers is summarized in the following function 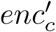 shown below. It is formulated recursively where in the *i*-th step of the recursion, substring *x*[*i, k* − *i*+1] is processed. The encoding of a palindromic *k*-mer solely consists of unspecific pairs until reaching the middle of the *k*-mer, which is either the empty string (even *k*) or a single character (odd *k*), the latter of which is encoded by its rank. For unspecific pairs and the remainder, the order of digits in the encoding corresponds to the order of characters in the given sequence. In contrast, for a reason explained later, the two digits encoding a specifying pair are placed next to each other at positions 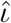 and 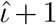 as shown in Example 1.

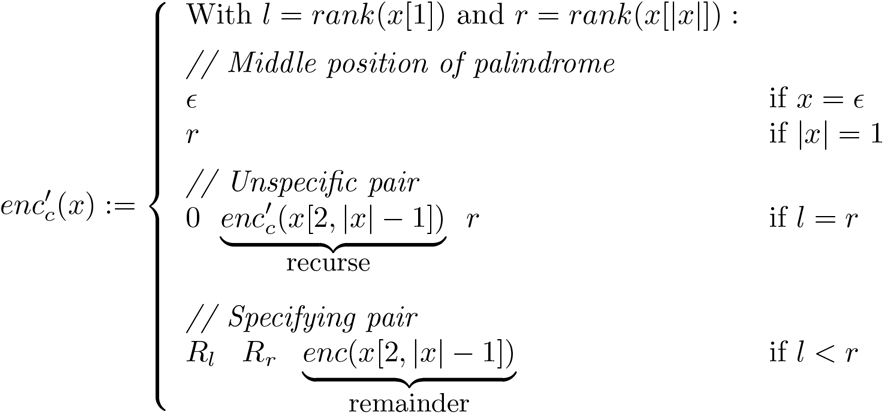

##### Example 1.

*Let k* = 6 *and σ* = 4. *The k-mer x shown below contains one unspecific pair. The specifying case is highlighted in bold and the remainder is underlined. R*(0, 1) = 0 *such that R*_*l*_ = 1 *and R*_*r*_ = 0 *(see Table 1)*.

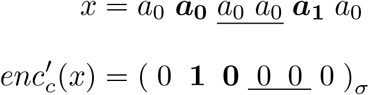

### 3.2 Transformation to a minimal encoding

In 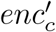, for a specifying pair, not all combinations of digits are used. Thus, the image of 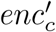 is not an interval. For a given 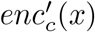, however, knowing the position 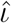 of the specifying pair (if *x* is non-palindromic), we can calculate 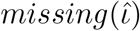, the number of all smaller elements of the co-domain that are not an encoding of any canonical *k*-mer. By subtracting this number from the decimal value of 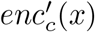, we obtain a continuous range of encodings from 0 to 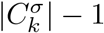 and thus a minimal encoding.

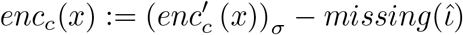

To determine the value of 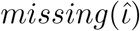, we need to count the number of digit sequences that (i) are not a valid encoding by 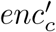 and (ii) have a value smaller than 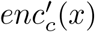. For palindromes, no such sequences exist, i.e., 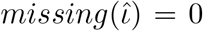. Otherwise, regarding (i), a sequence of digits can be identified invalid by reading from left to right and observing neither a zero that would result from encoding an unspecific pair, nor any of the valid 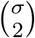 different combinations resulting from encoding a specifying pair. There are 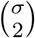 such invalid combinations of digits *a*,*b* with 0 *< a < b*. Regarding (ii), observe that in 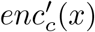, to the left of position 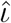 there are only zeros. We have to consider those sequences starting with further zeros, i.e., positions of invalid pairs *j, j* + 1 that are to the right of 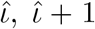. Further observe that any such pair has to be left of the middle position of the *k*-mer, i.e., 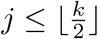. For each such pair *j, j* + 1, the remaining *m* := *k* − (*j* + 1) positions can take any value such that *σ*^*m*^ many invalid sequences have to be subtracted each.

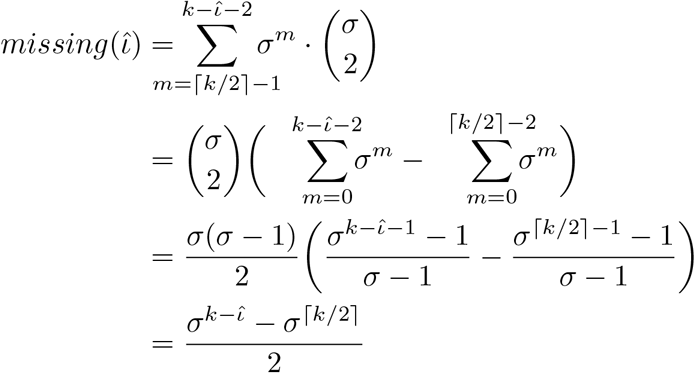

#### Example 2.

*Consider an alphabet of size σ* = 4, *k-mer length* 6, *and k-mer x* = *a*_0_ *a*_0_ *a*_0_ *a*_0_ *a*_1_ *a*_0_ *from Example 1*. *In this case*, 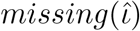 *counts all elements of the co-domain that are not in the image and have a value smaller than* 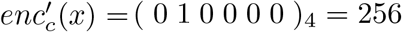. *Those are all sequences of the form*

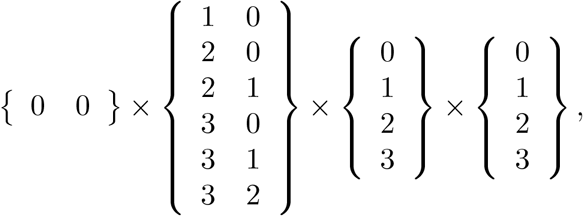

*i.e*., *they begin with two zeros, followed by a pair of digits that does not encode a specifying pair, followed by any two remaining digits. The number of these combinations is* 6 · 4 · 4 = 96, *i.e*., 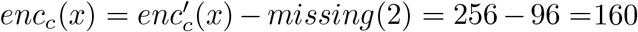. *The next lowest valid case is: enc*_*c*_(*a*_3_ *a*_3_ *a*_2_ *a*_3_ *a*_3_ *a*_3_) = ( 0 0 2 1 3 3 )_4_ − *missing*(3) = 159 − 0 = 159.

#### Theorem 1.

*Function enc*_*c*_ *is a minimal encoding for* 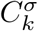.

*Proof*. First observe that 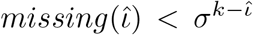, and thus position 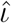 in 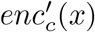 will not be affected by subtracting 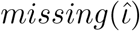 and, in particular, the number of leading zeros is the same in 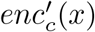 and *enc*_*c*_(*x*). Further observe that the number of leading zeros is equal to 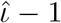, and thus allows us to derive 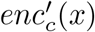 from *enc*_*c*_(*x*) by adding 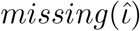. Hence, there is a bijection between the images of *enc*_*c*_ and 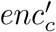.

A careful inspection of 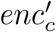 shows that it is well defined on any canonical *k*-mer and 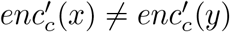 for any *x ≠ y*: the number of leading zeros of 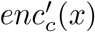 together with the corresponding digits at its end determine the unspecific pairs. Either the following two digits determine the specifying pair and the remaining digits determine the remaining characters, or the middle position is reached in case of a palindrome.

It remains to be shown that the 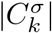 many different canonical *k*-mers are mapped to values not larger than 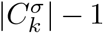. To this end, consider a value larger than 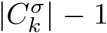 to be decoded. In base-*σ* space, this value would take at least 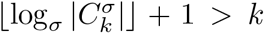 digits. Since valid encodings are of length *k* (including leading zeros), this is a contradiction. □

This encoding can easily be extended to include non-canonical *k*-mers such that they are encoded as their canonical counterpart. To this end, the specifying cases have to be complemented with those pairs that specify a string being non-canonical, i.e., *R*(*a, b*) := *R*(*b, a*) for *b < a*. Then, instead of the remainder, its reverse has to be encoded.

### 3.3 Palindromes

Palindromes do not pair to one canonical *k*-mer but are usually expected to be observed in sequence data approximately as often as non-palindromic *k*-mers that *do* pair to one canonical *k*-mer. Thus, palindromes might require special consideration in some applications, e.g., in statistical analyses, or for an even distribution of observed *k*-mers into buckets for parallel processing.

The above encoding allows a simple differentiation of palindromes from other *k*-mers. It is designed in such a way that palindromes are mapped to the lowest *σ*^⌈*k/*2⌉^ values, which can be seen as follows. If and only if *x* is a palindrome, 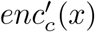 has ⌊*k/*2⌋ leading zeros. Otherwise, the number of leading zeros is smaller. As mentioned in the proof of Theorem 1, the number of leading zeros is the same for *enc*_*c*_(*x*). Thus, the encoding of palindromes has more leading zeros than any other encoding.

#### Example 3.

*Consider an alphabet of size σ* = 4 *and k-mer length* 2. *In this case*, 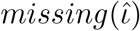 *is always zero such that enc* = *enc*^′^. *All k-mers in order of their encoding are shown below*.

| encoding | $k\text{-mer}(s)$ | |
| --- | --- | --- |
| 00 | $a_0a_0$ | } *palindromes* |
| 01 | $a_1a_1$ | |
| 02 | $a_2a_2$ | |
| 03 | $a_3a_3$ | |
| 10 | $a_0a_1 = (a_1a_0)^{-1}$ | |
| 11 | $a_0a_2 = (a_2a_0)^{-1}$ | |
| 12 | $a_0a_3 = (a_3a_0)^{-1}$ | |
| 13 | $a_1a_2 = (a_2a_1)^{-1}$ | |
| 20 | $a_1a_3 = (a_3a_1)^{-1}$ | |
| 21 | $a_2a_3 = (a_3a_2)^{-1}$ | |

This characteristic of *enc*_*c*_, mapping palindromes to the smallest values, will also be exploited in the following section, where palindromes play a particular role.

## 4 Canonicalization under reverse complementation

In certain applications, in particular when handling genomic sequences, instead of only reverting a sequence, its reverse (Watson-Crick) complement is considered. For an even-size alphabet {*a*_0_, … , *a*_*σ*−1_}, the *reverse complement* of a string *s* is 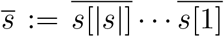, where 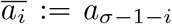. For instance, for the DNA alphabet 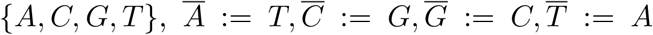. An extension to alphabets of odd size, where 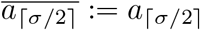 would be possible, but those technicalities are omitted here for the sake of comprehensibility. Let 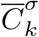 be the set of all canonical *k*-mers over an alphabet of even size *σ* under reverse complementation.

For an odd-length *k*-mer, taking its reverse complement always changes the character at its middle position. Thus, *x* is always different from 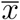, i.e., there are no palindromes, and the number of canonical *k*-mers is half the number of *k*-mers.

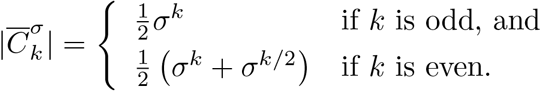

The encoding 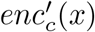 from Section 3.1 can be applied analogously here – just 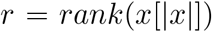 has to be replaced by *r* = *rank*(*x*[|*x*|]), which we will denote 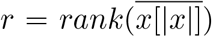 in the following. If *k* is odd, there are no palindromes, but the middle position of a *k*-mer might specify it being canonical. So, the very first case in the definition of 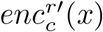 will only apply to one half of the alphabet. Hence, the lowest *σ/*2 · *σ*^⌊*k/*2⌋^ = 1*/*2 · *σ*^⌈*k/*2⌉^ values are never taken (see also Section 3.3). Their number has to be included in 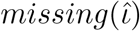 such that the image is shifted from 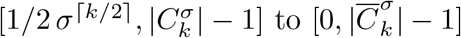.

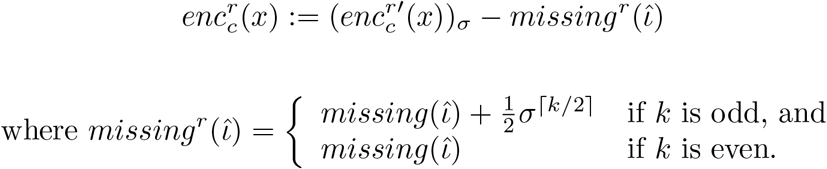

### Example 4.

*Let k* = 5 *and σ* = 4. *The k-mer y shown below contains two unspecific pairs. It is specified being canonical by its middle position*, 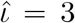, *highlighted in bold*.

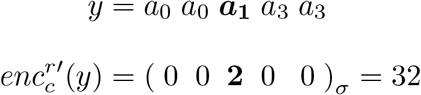

*There is no k-mer with a smaller encoding, because this would require a middle position encoded with a value lower than 2, i.e. character a*_2_ *or a*_3_, *which contradict canonicity. We have* 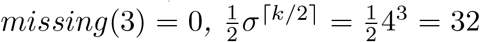, *and thus missing*^*r*^(3) = 32 *and* 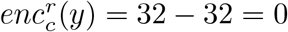.

### Corollary 1.

*Function* 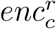 *is a minimal encoding for* 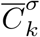.

To allow non-canonical *k*-mers as input to be encoded as their canonical counterpart, instead of the specifying pair and the remainder, their reverse complements have to be encoded.

## 5 Efficient bit-based implementation for the DNA-alphabet

Here, we consider the commonly used DNA alphabet {*A, C, G, T*} with 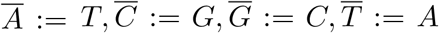. For the sake of convenience, we consider *k* being odd such that palindromes can be left aside in the following explanations and the number of canonical *k*-mers is 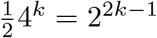.

### 5.1 Space efficiency

The four different rank values can be binary encoded by two bits each: *rank*(*A*) = 0 = (00)_2_, *rank*(*C*) = 1 = (01)_2_, *rank*(*G*) = 2 = (10)_2_ and *rank*(*T* ) = 3 = (11)_2_. Since both the standard rank based encoding *enc* and the encoding for canonical *k*-mers 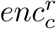 result in *k* digits, any encoding is of length 2*k* bit. However, the encodings of 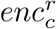 only span the integer range from 0 to 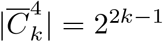 such that only log_2_(2^2*k*−1^) = 2*k* − 1 bits are necessary. This means, the first bit of any 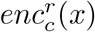 will always be zero and can thus be omitted to store any canonical *k*-mer using 2*k* − 1 bits.

### 5.2 Time efficiency

When all overlapping *k*-mers of a given (long) string *s* have to be encoded, we want to avoid processing each single character *k* times: for all *k*-mers it is contained in. Here, we provide an algorithm to compute the encoding of a *k*-mer *s*[*i*..*i* + *k* −1] from the encoding of the previously read *k*-mer *s*[*i*−1..*i* + *k*] in constant time, i.e., with a run time complexity independent of *k*, assuming *k*-mers fit into a constant number of computer words.

For the standard rank based encoding *enc* in 2-bit representation, such a rolling hash function can easily be formulated: drop the two bits corresponding to *s*[*i* −1], shift *enc*(*s*[*i*−1..*i*+*k*]) by two bits to the left, and insert *enc*(*s*[*i*+*k* −1]) corresponding to the new last character from the right.

For the minimal encoding 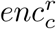, a shift/update approach is not applicable, because, within a *k*-mer *x*, the encoding of a character *x*[*i*] depends on its mate *x*[*k* − *i*+1], and this pairing changes with each shift. Instead, since 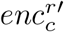 encodes many characters simply by their rank, we maintain a standard encoding (in 2-bit representation) of the current *k*-mer *x* as well as 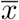 using the constant-time shift/update function described above and modify (a copy of) *enc*(*x*) to obtain 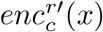 using a constant number of atomic operations and finally subtract 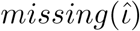.

Let *p* denote the length of the longest common prefix of *x* and 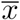, i.e., the number of unspecific pairs in *x*. Note that *p < k/*2. This value can be computed in constant time by performing an *exclusive-or* operation on *enc*(*x*) and 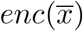. Since the binary encoding of a character is the negation of its complement, unspecific pairs will result in a prefix of zeros, the length of which can be obtained in constant time by the operation *count leading zeros (clz)*, which (or an equivalent operation of which) belongs to the repertoire of atomic CPU instructions on modern architectures. An integer division by two yields *p*.

Recall that from left to right, the encoding is composed of the following four elements:

1. a prefix of 2*p* zero-bits for the characters to the left of the specifying case,
2. four bits for the specifying pair (or two bits for a specifying middle character),
3. the encoded remainder (characters within the specifying pair, if any),
4. the encoded characters to the right of the specifying case (if any).

In the given standard 2-bit encoding of a *k*-mer, all elements can be localized with respect to *p*. Elements 3 and 4 can be extracted using bit masks and a constant number of *and* operations. Element 3 has to shifted by 2 bits, and element 4 has to be complemented, which can be done using an *xor 1* operation. Bits for element 2 can be replaced according to 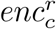 with a constant number of operations. It remains element 1, i.e., replacing a prefix of length 2*p* bit by a sequence of zeros. This can also be performed in constant time by using an *and* operation with a corresponding bit mask of 2*p* zeros followed by ones. All necessary bit masks for ⌊*k/*2⌋ putative values of *p* can be precomputed.

#### Example 5.

*Consider k* = 6 *and the following k-mer z, where the specifying pair is highlighted in bold and the remainder is underlined. R*(*rank*(*C*), *rank*(*C*)) = *R*(1, 2) = 3 *such that R*_*l*_ = 1 *and R*_*r*_ = 3 *(see Table 1)*.

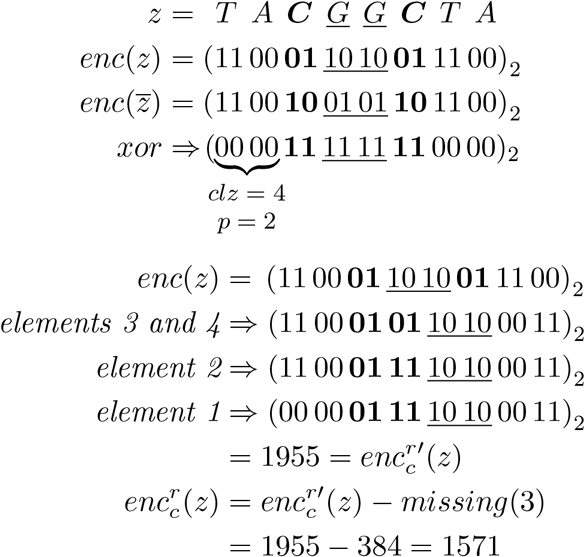

The above argumentation can easily be extended to even-length *k*-mers and yields the following lemma.

#### Lemma 1.

*For a k-mer x, given the encodings enc*(*x*) *and* 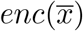, *the encoding* 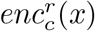 *can be computed in constant time, assuming* 2*k bits fit into a constant number of computer words*.

Once the first *k*-mer of a string and its reverse complement have been 2-bit encoded, this and all successive *k*-mers can be encoded by 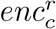in time linear in the sequence length and independent from *k*.

#### Corollary 2.

*All k-mers of a sequence s can be encoded by the minimal encoding enc*_*c*_ *in O*(*k* +|*s*|) *time, assuming* 2*k bits fit into a constant number of computer words*.

An exemplary implementation of this approach can be found under https://gitlab.ub.uni-bielefeld.de/gi/MinEncCanKmer.

## 6 Conclusions

We proposed a general minimal encoding for canonical *k*-mers of arbitrary (even or odd) length on alphabets of arbitrary size. The approach applies to both cases: considering reverse strings or reverse complements.

Canonical *k*-mers are not evenly distributed within a standard rank based encoding. In particular, *k*-mers on the DNA alphabet are not evenly distributed within a standard 2-bit encoding. The presented encoding maps all *n* canonical *k*-mers to a continuous interval of integers from 0 through *n* −1 (it is *minimal* ). Furthermore, our encoding has the additional property that palindromic *k*-mers, which are statistically observed more often than other canonical *k*-mers and thus require special consideration, are mapped to a specific range of integers.

For the prevailing case of *k*-mers on the DNA alphabet, our approach allows to encode all *k*-mers of a sequence *s* in *O*(*k* + |*s*|) time using a shift/update approach and efficient bit operations.

In contrast to non-surjective hash functions, a minimal encoding renders the possibility of using array-based data structures for efficiently storing canonical *k*-mers. Further, since the encoding (of odd length *k*-mers) requires one bit fewer than a standard 2-bit encoding, it can be of use to save space when canonical *k*-mers are not stored in individual computer words (of even length), e.g. in [10].

Even though substantial contributions on the practical side are not evident, this work contributes with basic research in the field of *k*-mers by closing obvious theoretical gaps.

Another line of research would be efficiently encoding *homopolymer compressed k*-mers, which are *k*-mers in which runs of the same characters are replaced by a single copy of the characters each. This restricts the set of *k*-mers under consideration to 4 · 3^*k*−1^ and thus, again, raising the question for a minimal encoding.

## Acknowledgments

I thank Fabian Kolesch for pointing me to the problem, the DSB community (in particular Antoine Limasset and Sven Rahmann) for input and motivation, and the reviewers for substantial feedback.

## Footnotes

1 https://cs.stackexchange.com/questions/82644/compact-mapping-from-an-involuted-set (Thanks to Antoine Limasset for this pointer.)

